# Cancer cell lipid class homeostasis is altered under nutrient-deprivation but stable under hypoxia

**DOI:** 10.1101/382457

**Authors:** Jan Lisec, Carsten Jaeger, Nousheen Zaidi

**Author notes:** **Corresponding author**: Nousheen Zaidi.

## Abstract

Under oxygen/nutrient deprivation cancer cells modify the balance between fatty acid (FA) synthesis and uptake, which alters the levels of individual triglyceride or phospholipid sub-species. These modifications may affect survival and drug-uptake in cancer cells. Here, we aimed to attain a more holistic overview of the lipidomic profiles of cancer cells under stress and assess the changes in major lipid-classes. First, expressions of markers of FA synthesis/uptake in cancer cells were assessed and found to be differentially regulated under metabolic stress. Next, we performed a broad lipidomics assay, comprising 244 lipids from six major classes, which allowed us to investigate robust stress induced changes in median levels of different lipid classes -additionally stratified by fatty acid side chain saturation status. The lipidomic profiles of cancer cells were predominantly affected by nutrient-deprivation. Neutral lipid compositions were markedly modified under serum-deprivation and, strikingly, the cellular level of triglyceride subspecies decreased with increasing number of double bonds in their fatty acyl chains. In contrast, cancer cells maintained lipid class homeostasis under hypoxic stress. We conclude that although the levels of individual lipid moieties alter under hypoxia, the robust averages of broader lipid class remain unchanged.

## Introduction

Lipid metabolism has emerged as an important aspect of cancer cell metabolism and is widely shown to be associated with various malignant processes (1-3). Cancer cells require a constant supply of lipids for membrane biogenesis and protein modifications. Several studies have shown that, in order to cope with these increased demands, cancer cells activate *de novo* lipid synthesis pathways (4-8). Fatty acid synthase (FASN) –a key-regulator of *de novo* fatty acid (FA) synthesis– has been extensively shown to fuel cancer cell proliferation and malignant progression (3). Expression of 3-hydroxy-3-methylglutaryl-CoA reductase (HMGCR) –the rate-controlling enzyme of the mevalonate pathway– is also up-regulated in cancers (9). Importantly, inhibition of FA synthesis or cholesterol synthesis pathways results in growth-arrest of lipogenic tumor cells rendering these pathways interesting targets for antineoplastic therapy (4, 10-16). Although endogenous FA synthesis has historically been considered the principal source of fatty acids (FAs) in cancer cells, lipolytic phenotypes are also widely recognized (reviewed in (17)). For example, it has been reported that in addition to the markers of *de novo* synthesis (FASN) different cancer cells also express markers of lipolysis (lipoprotein lipase, LPL) and exogenous FA uptake (CD36) (18). Additional support for coordinated lipolytic and lipogenic metabolism in cancer cells involves the incorporation of endogenously synthesized FAs into cellular neutral lipid stores. Nomura *et al*. (19) have proposed that complementary lipolytic pathways are required to release fatty acyl moieties from these lipid reservoirs and have demonstrated a specific role for an intracellular lipase, monoacyl glycerol lipase (MGLL), in promoting tumorigenesis. MGLL provides, by de-esterification, a stream of intracellular free FAs to fuel proliferation, growth, and migration. Taken together, these findings are compatible with the notion that both lipogenesis and lipolysis may be utilized by cancer cells to fulfill their FA requirements.

The mode of FA acquisition –via *de novo* synthesis or uptake– may significantly affect the lipidomic profiles of cancer cells. Mammalian cells have a limited ability to synthesize polyunsaturated fatty acids *de novo*, as they lack the Δ12 desaturase. Therefore, enhanced *de novo* FA synthesis enriches the cancer cell membranes with saturated and/or mono-unsaturated fatty acids (20). As these FAs are less prone to lipid peroxidation than polyunsaturated acyl chains, *de novo* synthesis was proposed to make cancer cells more resistant to oxidative stress-induced cell death (20). Moreover, as saturated lipids pack more densely their increased levels alter lateral and transverse membrane dynamics that may limit the uptake of drugs, making the cancer cells more resistant to therapy (20). Hence, the balance between FA synthesis and uptake may have important therapeutic implications.

Cancer cells are shown to modify balances of FA synthesis and uptake under metabolic stress, i.e. induced by oxygen and nutrient deprivation (3, 21). This metabolic flexibility is particularly important for cancer cells within solid tumors that are exposed to oxygen- and nutrient-gradients depending on their distance from the nearest blood vessels. The inefficient vascularization limits access of various nutrients – such as amino acids, sugars and lipids– to tumor tissues. The effect of oxygen/nutrient deprivation on leukemic cells was presumed to be inconsequential and remained long overlooked. This view is also being revised and hypoxia has been shown to influence leukemic cell proliferation, differentiation, and resistance to chemotherapy (reviewed in (22)). It will also be interesting to study the impact of oxygen/nutrient deprivation on metabolic pathways in leukemic cells.

To date, the effect of metabolic stress on lipid metabolism in cancer cells appears inconsistent. For example, under hypoxia the expression of FASN is reported to be significantly up-regulated in breast cancer cells (23), down-regulated in HepG2 liver cancer cells (24) and unaffected in HCT116 colorectal cancer cells (25). Certain types of cancer cells are shown to switch from lipid synthesis to uptake under hypoxia. Multiple studies indicate that under hypoxic conditions cancer cells increase FA uptake (21, 26), particularly of lipid species containing monounsaturated acyl chains (21). Most studies on the impact of hypoxia on lipidomic profiles of cancer cells are limited to specific lipid classes. For instance, it was shown that under hypoxia breast cancer cells display modified phospholipid profiles mainly characterized by the presence of shorter and more saturated acyl chains while other lipid classes were not considered (27). Another study reported that under hypoxic conditions cellular levels of triglycerides with three double bonds were significantly decreased in MCF7 breast cancer cells (26), but significantly increased in U87 glioblastoma cells (26), but data on membrane lipids were not collected. Nutrient deprivation, specifically low-lipid environment, is also shown to affect *de novo* lipid synthesis pathways in cancer cells (28, 29). It has been reported that cancer cells from different origins differentially activate and thrive on *de novo* lipid synthesis pathways in a low-lipid environment. It was shown that cancer cells that attain the highest lipogenic activity under lipid-reduced environment are best able to cope with lipid reduction in term of proliferative capacity (28). Hence, the changes in lipid metabolism and lipidomic profiles of cancer cells that are induced by environmental stress appear to be cell-type specific. Together, these data indicate that both genetic background and environmental conditions may determine the relative dependence of cancer cells on *de novo* lipid synthesis versus lipid uptake.

Herein, to study the complex interplay between metabolic stress and lipid metabolism in cancer cells, we selected a biologically diverse panel of cancer cell lines –three leukemia cell lines (to cover intra-group variance), two colon cancer cell lines and one lung cancer cell line. We were mainly interested in studying the impact of physiologically relevant metabolic stress on lipid metabolism in cancer cells. To achieve that cancer cells were cultivated under nutrient-deprivation and hypoxia –that closely mimics the *in vivo* conditions. The expression of relevant markers was assessed to study the relative dependence of cancer cells on *de novo* lipid synthesis versus lipid uptake/degradation pathways under metabolic stress. In order to gain more systematic insight on the effects of metabolic stress on lipidomic profiles we performed a broad lipidomics assay comprising 244 lipids from six major classes. To this end we identified multiple changes in lipidomic profiles of cancer cells cultivated under low-serum or lipid-deficient conditions. Under hypoxic stress cancer cells displayed alterations in cell proliferation rates and expression profiles of various lipid metabolism associated genes. Interestingly, no robust changes were observed in lipidomic profiles of hypoxic cancer cells indicating that the cells maintain lipid class homeostasis.

## Materials and Methods

### Cell culture and treatments

The SW480, SW620, A549, KG1, KCL22 and KU812 cell lines were maintained in DMEM (Gibco, 31966-021) or RPMI 1640 medium (Gibco, 61870-010) media supplemented with 10% fetal bovine serum (FBS) (Sigma, F75240) and penicillin-streptomycin solution (Corning, 30-002-CI). Cell cultures were maintained in the atmosphere of 5% CO_2_ and 37°C. For all experiments cells were initially seeded and cultivated in normal media for 24 hours. Then to induce metabolic stress media and/or growth conditions were respectively changed and cells were cultivated for additional 48 hours under either one of the following condition: lipoprotein deficient medium (LPDS serum), low-serum (LS) medium (2% serum), hypoxia (2% O2), or hypoxia in combination with LS medium. For lipoprotein deficient conditions the media were supplemented with lipoprotein deficient serum (LPDS) that was purchased from Merck (LP4) and used according to manufacturer’s guidelines. For determining the cells number cells were stained with trypan blue and counted using Countess^®^ automated cell counter (Invitrogen). Cell lines were commercially authenticated (Eurofins, Germany) and mycoplasma tested prior to submission of this manuscript.

### Quantitative RT-PCR

For quantitative RT-PCR, total RNA was extracted from cell pellets using Quick-RNA™ MiniPrep Plus (Zymo Research). All RNA samples were reverse-transcribed into cDNA using SuperScript™ III Reverse Transcriptase (Thermo Scientific, 18080093) and Oligo(dT)18 Primers (Thermo Scientific, SO131). Quantitative PCR was performed using a TaqMan™ Gene Expression Master Mix (4369016, Applied Biosystems) *via* StepOne Real-Time PCR Systems (Applied Biosystems). The TaqMan Gene Expression assays used were Hs01005622_m1 (fatty acid synthase, FASN), Hs00168352_m1 (3-hydroxy-3-methylglutaryl-CoA reductase, HMGCR), Hs00996004_m1 (monoglyceride lipase, MGLL), Hs00173425_m1 (lipoprotein lipase, LPL) and Hs00354519_m1 (CD36). The expression of each gene was normalized to the expression of GADPH (Hs02786624_g1).

### Lipid Extractions

First, the cell pellets were washed with 0.5 mL 0.9% NaCl. For extraction of lipids the pellets were resuspended in 1 ml ice-cold MMC (1:1:1 v/v/v methanol/MTBE/chloroform). Samples were incubated on an ultrasonic bath for two minutes. Phase separation was induced by adding 300 μL MS-grade water. After 10 min incubation, the samples were centrifuged for 10 min at 1000 rpm and the upper (organic) phase was collected. Then 200 μL of collected organic phase were dried in a vacuum rotator and stored at –20 °C until analysis.

### Lipidomic Profiling

Dried sample extracts were reconstituted in 100 µL 2:1:1 v/v/v isopropanol/acetonitrile/water. 5 µL aliquots were injected into an ACQUITY I-class ultra-performance liquid chromatography (UPLC) system (Waters, Germany) coupled to an Impact II high-resolution quadrupole time-of-flight mass spectrometer (Bruker Daltonik GmbH, Germany). Chromatographic separation was achieved by gradient elution (%A: 0 min, 60; 1.2 min, 57; 1.26 min, 50; 7.2 min, 46; 7.26 min, 30; 10.8 min, 0; 12.96 min, 0; 13.02 min, 60; 14.4 min, 60) using a buffered solvent system (A: 60:40 v/v acetonitrile/water, B: 90:10 v/v isopropanol:water, both with 10 mM ammonium formate and 0.1% formic acid) and a 2.1 mm × 75 mm × 1.7 µm CSH-C18 column (Waters, Germany) equipped with an 0.2 µm pre-column in-line filter. The flow rate was 0.5 mL min^-1^ and column temperature was 55°C. Electrospray ionization (ESI) conditions were as follows: polarity (+), capillary voltage, 4500 V, end plate offset, 500 V, nebulizer pressure, 2.5 bar, dry gas (N2) flow, 8 L/min. Ion transfer parameters were set to: Funnel 1 RF, 200 Vpp, Funnel 2 RF, 200 Vpp, Hexapole RF, 50 Vpp, Quadrupole Ion Energy, 5 eV, Low Mass, 100 m/z, Collision Energy, 8.0 eV, Pre Pulse Storage, 6.0 µs, Stepping Mode, Basic, Collision RF, 500-1000 Vpp, Transfer Time, 60-100 µs, Timing, 50/50, Collision Energy, 100-250%. Alternating MS and MS/MS scans were acquired using a Sequential Windowed Acquisition of All Theoretical Fragment Ion Mass Spectra (SWATH) scheme (m/z 350-975, width 25 Da). For internal calibration, Na formate clusters were spiked into the LC effluent at the end of each run.

After data acquisition, files were converted to Analysis Base File (ABF) format using a publicly available converter (Reifycs, Japan) and imported into MS-DIAL (Tsugawa et al. 2015). MS-DIAL parameter settings were as follows: Soft Ionization, Data independent MS/MS, Centroid data, Positive ion mode, Lipidomics. Detailed analysis settings were left at default, except for Retention time end (10 min), Alignment Retention time tolerance (0.2 min), Identification Retention time tolerance (3 min) and Identification score cut off (60%). Identified peaks were exported to a text file and subjected to statistical analysis.

### Statistical analysis

The differences between groups were analyzed by ANOVA or t-test (paired or unpaired), where applicable. Statistical analyses and graphical representations for lipidomic data and quantitative RT-PCR data were performed using the R software environment 3.4.2 (http://cran.r-project.org/) or MetaboAnalyst 3.5 (http://www.metaboanalyst.ca/faces/home.xhtml). P-values <0.05 were considered statistically significant and indicated when different.

## Results

### Comparison of baseline expression of selected markers for *de novo* lipid synthesis, lipid uptake/degradation in cancer cell lines

We first compared the baseline expression of the key genes involved in *de novo* lipid synthesis (FASN and HMGCR) in a biologically diverse panel of cancer cell lines (KG-1, KCL22, KU812, SW480, SW620 and A549). KG1 is an acute myeloid leukemia cell line, whereas KCL22 and KU812 are chronic myeloid leukemia cell lines. SW480 and SW620 are colon carcinoma cell lines derived from primary tumor and lymph-node metastasis of the same patient, respectively. A549 is a human lung carcinoma cell line We observed that the expression levels of FASN –key regulator of *de novo* fatty acid synthesis–significantly varied across the selected cell lines with up to ~40-folds difference between KG1 and SW480 cells (**Figure 1a, Supplementary Table 1**). The expression of HMGCR –rate-controlling enzyme of the mevalonate pathway– also varied significantly among the selected cell lines with up to ~42-folds difference between KG1 and SW480 (**Figure 1a, Supplementary Table 1**). The variance in expression patterns of HMGCR was similar to that of FASN. Regression analysis revealed highly positive correlation (R^2^=0.86; P <0.0001) between baseline mRNA levels of FASN and HMGCR (**Supplementary Figure 1**). All the leukemia cell lines had higher expression of both FASN and HMGCR in comparison to the adherent cell lines derived from solid tumor tissues. It reflects comparatively higher *de novo* lipid synthesis in leukemia cells.

**Figure 1:**
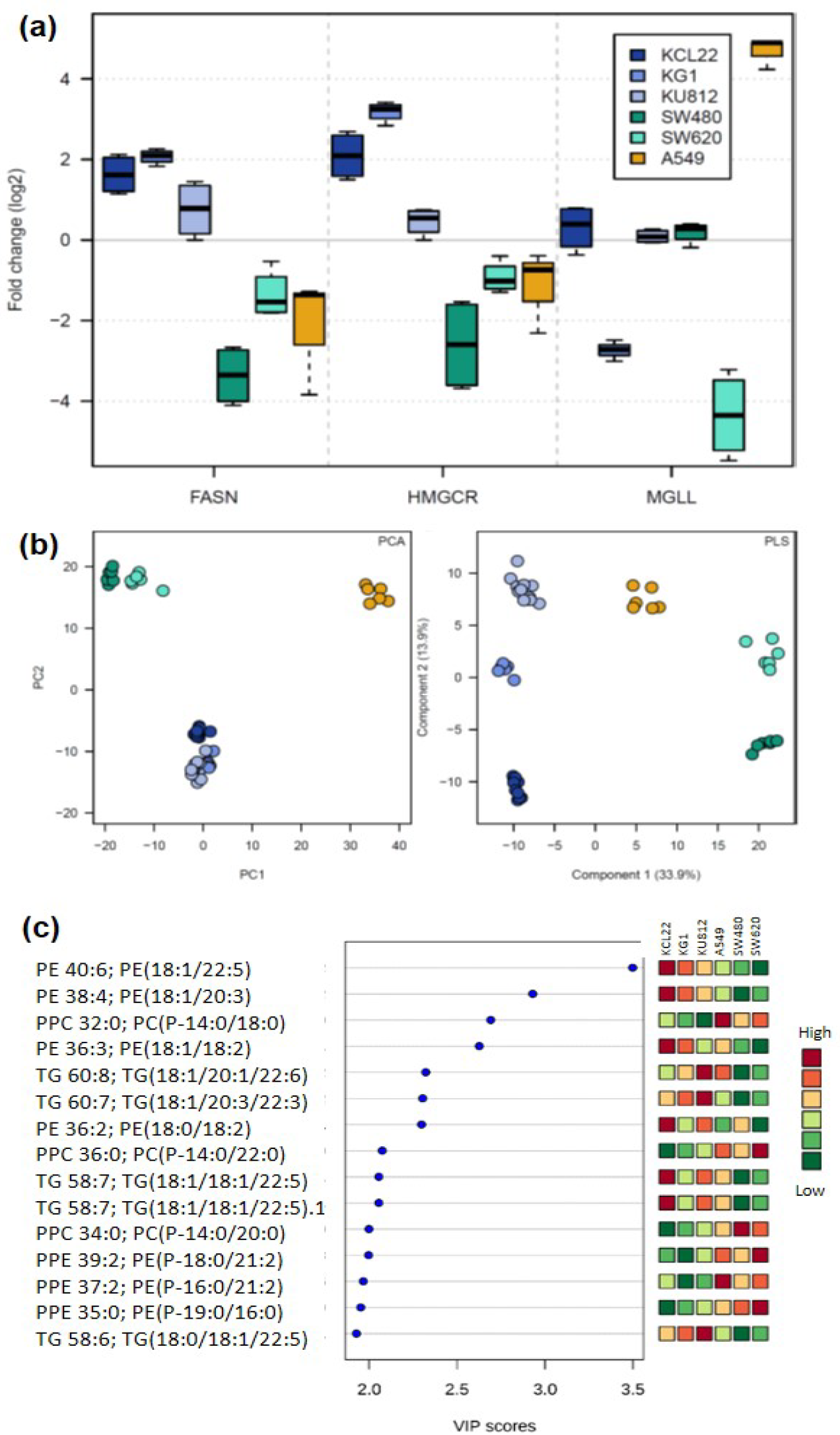
Baseline expression of various lipid metabolism-associated genes and lipidomic profiles in selected cancer cell lines. **(a)** Box plots showing log2 transformed values for baseline expression levels of FASN, HMGCR and MGLL in KG1, KCL22, KU812, SW480, SW620 and A549. The levels of the indicated transcript were measured in 3 to 6 samples by qPCR. The results show the distribution of indicated transcript relative to GAPDH and normalized to the median expression level of the corresponding transcript across all cell lines, with the box indicating the 25th–75th percentiles, with the median indicated. The whiskers show the range. Statistical significance was tested with ANOVA analysis followed by post-hoc Tukey’s test (see supplementary table 1). **(b)** *(Left-panel)* Principal component analysis (PCA) of lipidomic profiles KG1, KCL22, KU812, SW480, SW620 and A549 cell lines at baseline level. Percentage of the variance captured by each principal component (PC) is given close to each respective axis. *(Right-panel)* PLSDA model analysis of 244 common lipid molecules to differentiate six different cell lines (i.e. KG1, KCL22, KU812, SW480, SW620 and A549) **(c)** Potential discriminatory lipid molecules identified through VIP scores (VIP values of >2.0) derived from PLS-DA modeling of complete data matrix. Resulting VIP scores for top 15 lipid molecules are shown in increasing order of VIP score values to highlight their discriminatory potential.

The expression of lipolytic markers (LPL and MGLL) was also assessed for the selected cell line panel. We observed consistent expression of LPL only in SW480 and KU812. The expression of MGLL was strikingly different among the selected cell lines with up to ~250-folds difference between SW620 and A549 (**Figure 1a)**. Expression of CD36 –fatty acid uptake marker– was only detectable in KU812. These data indicate that the selected cell lines are significantly different in terms of baseline levels of lipid metabolism related transcripts.

### Comparison of baseline lipidomic profiles of cancer cell lines

Next, we compared the baseline lipidomic profiles of the selected cell lines. To integrate the lipidomics data with gene expression profiles we determined the lipidomic profiles of the same samples that were used for qPCR analysis. Global lipidomic profiling using Liquid Chromatography-Mass Spectrometry (LC-MS) followed by automatic annotation using MS-Dial (v.2.72 (30)) allowed to detect 244 lipid compounds each present in at least 90% of all samples (**Table 1**). Majority of the detected lipid molecules were either phospholipids (n=136) or neutral lipids (n=97). The major phospholipid subclasses comprised Phosphatidylcholines (PC, n=45), Phosphatidylethanolamines (PE, n=43), Plasmenylphosphatidylethanolamines (PPE, n=29) and Plasmenylphosphatidylcholines (PPC, n=8). The neutral lipids included subclasses of Triacylglycerols (TG, n=70), Cholesterol esters (CE, n=19) and Diacylglycerols (DG, n=7). Inherent differences in the lipidomic profiles were visualized using descriptive Principal Component Analysis (PCA) that showed a clear separation between different cancer cell lines (**Figure 1b**). As expected, cell line lipid profiles of similar tissue of origin are more similar. To further elucidate the individual compounds contributing predominantly to the variance observed in lipidomic data from the selected cell lines a partial least-square-discrimination analysis (PLS-DA) model was constructed. The PLS-DA score plots also showed clear separation between different cell lines (**Figure 1b**). Variable importance in the projection (VIP) values were applied (VIP values >2.0) to identify fifteen most important lipid molecules which mainly contributed in differentiating the lipidomic profiles of the cell lines (**Figure 1c**). Ten out of these lipid molecules were phospholipids, whereas five were triglycerides.

**Table 1.**

| **Table 1** |  |  |
| --- | --- | --- |
| **Lipid Type** | **Lipid Class** | **Number of molecules** |
| **Phospholipids** | Phosphatidylcholines (PC) | **45** |
|  | Lysophosphatidylcholines (lysoPC) | **2** |
|  | Lysophosphatidylethanolamines (lysoPE) | **4** |
|  | Phosphatidylglycerols (PG) | **1** |
|  | Plasmenylphosphatidylethanolamines (PPE) | **29** |
|  | Plasmenylphosphatidylcholines (PPC) | **8** |
|  | Phosphatidylethanolamines (PE) | **43** |
|  | Bis(monoacylglycero)phosphate (BMP) | **4** |
| **Neutral Lipids** | Cholesterol ester (CE) | **19** |
|  | Monoacylglycerol (MG) | **1** |
|  | Diacylglycerol (DG) | **7** |
|  | Triacylglycerol (TG) | **70** |
| **Sphingomyelins** | Sphingomyelins (SM) | **4** |
| **Acylcarnitine** | Acylcarnitine | **5** |
| **Lanosteryl oleate** | Lanosteryl oleate | **1** |
| **Lanosteryl palmitoleate** | Lanosteryl palmitoleate | **1** |
| **Total** |  | **244** |

Baseline lipidomic profiles for the selected cell lines were further analyzed accounting for saturation status (i.e. the largest number of double bonds in any acyl side chain). To this end, we only included lipid sub-groups with n>6 to ensure statistical robustness (CEs, DGs, PCs PPCs, PEs and TGs). To further increase robustness of the analysis, for each of the analyzed lipid classes all subspecies containing similar saturation status were averaged. **Figure 2** compares the levels of various lipids sup-species in the selected cell lines. To account for differences in cell number and hence total amount used in analysis we expressed each lipid peak intensity relative to the median peak intensity of the sample. Thereby, we can compare the relative distribution of lipid classes within a cell over various cell types. The most significant trends observed in these analyses were that the colon cancer cell lines displayed significantly lower levels of TGs subspecies containing ≥3 double bonds in comparison to other cell lines. Moreover, the A549 cells displayed significantly lower levels of PEs containing ≥3 double bonds in comparison to other cell lines.

**Figure 2:**
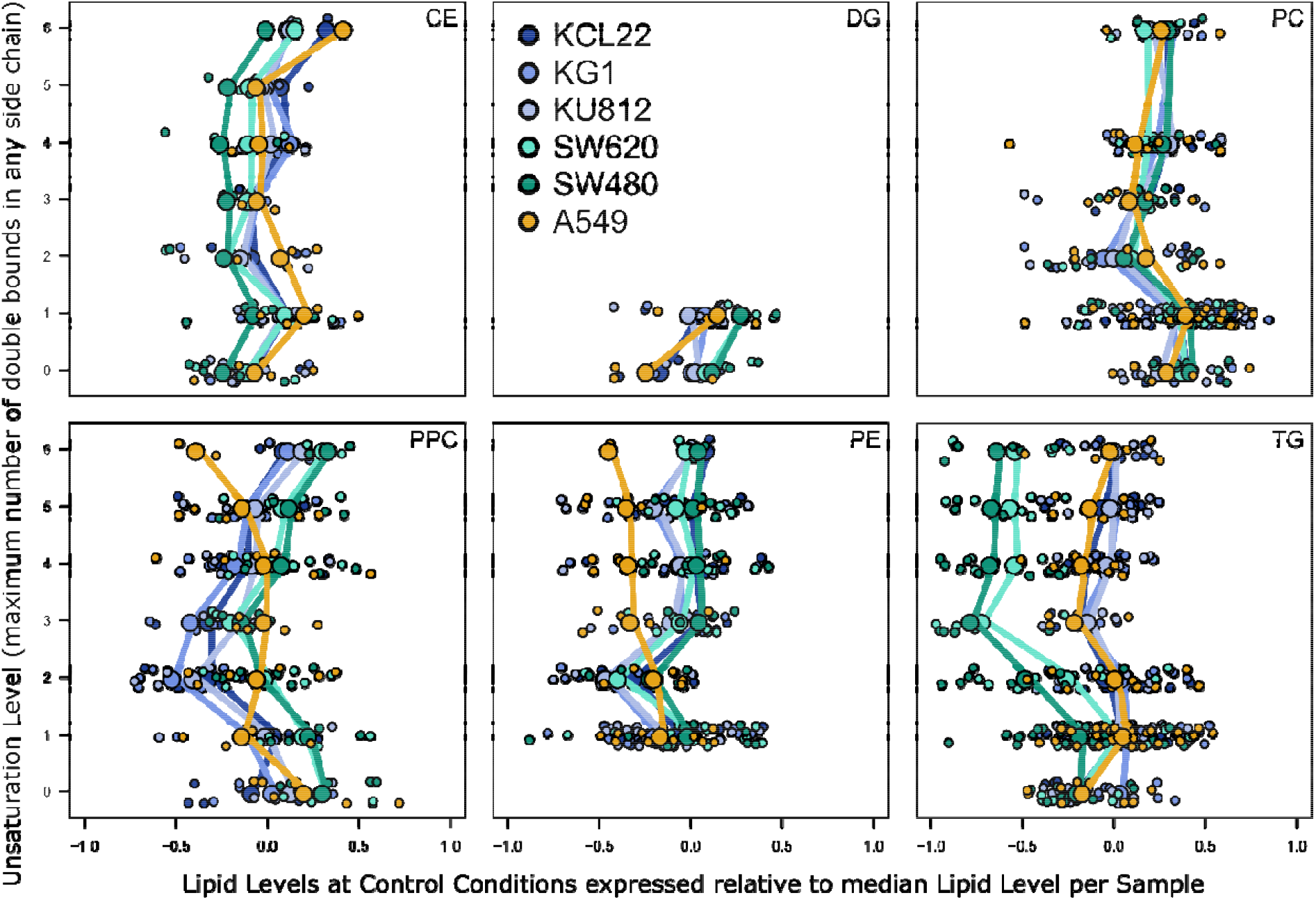
Baseline lipidomic profiles in selected cancer cell lines. Cellular levels of the six major lipid classes – including CEs, DGs, PCs PPCs, PEs and TG– were compared in KCL22, KG1, KU812, SW480, SW620 and A549 cells. Lipids are separated according to largest number of double bonds in any side chain observed (y-axis). Group means are depicted by larger symbols and connected with lines of similar color. Smaller symbols representing individual lipids were randomly scattered around saturation values (y-axis) to improve visibility. The data were log10-transformed and are expressed as/normalized to sample median (x axis).

### Effect of metabolic stress on cell proliferation and expression of different lipid metabolism-related genes in cancer cells

We have previously shown that cancer cells cultivated under low-lipid conditions display differential growth patterns (28). Here, we first studied the effect of different metabolically-stressed culture conditions on cell proliferation in various cancer cells. To induce metabolic stress, cells were cultivated for 48h under following conditions: lipoprotein deficient medium (LPDS serum), low-serum medium (2% serum), hypoxia (2% O2), or hypoxia in combination with low-serum medium. **Figure 3** shows proliferation rates of cancer cells cultivated under different cell culture conditions. Here, for each cell line the data were normalized to its proliferation rate under normal condition. Thereby, we remove the existing differences in baseline proliferation rates of selected cell lines (**Supplementary Figure 2**) and emphasize on stress-induced changes in proliferation rates. We observed that metabolic stress significantly impacted proliferation rates of most cell lines (**Figure 3**). KU812, SW480 and SW620 showed decreased proliferation rates when cultivated in media containing lipoprotein-deficient serum (LPDS). KU812 and SW480 express LPL and CD36 proteins that are involved in lipolysis and uptake of extracellular fatty acids. Therefore, they might be more sensitive to LPDS medium. All cell lines except SW620 showed reduced proliferation rates under low-serum environment. Hypoxic conditions induced decreased proliferation rates in A549 and SW480. When cultivated under hypoxia in combination with low-serum medium all cell lines except SW620 displayed reduced proliferation rates.

**Figure 3:**
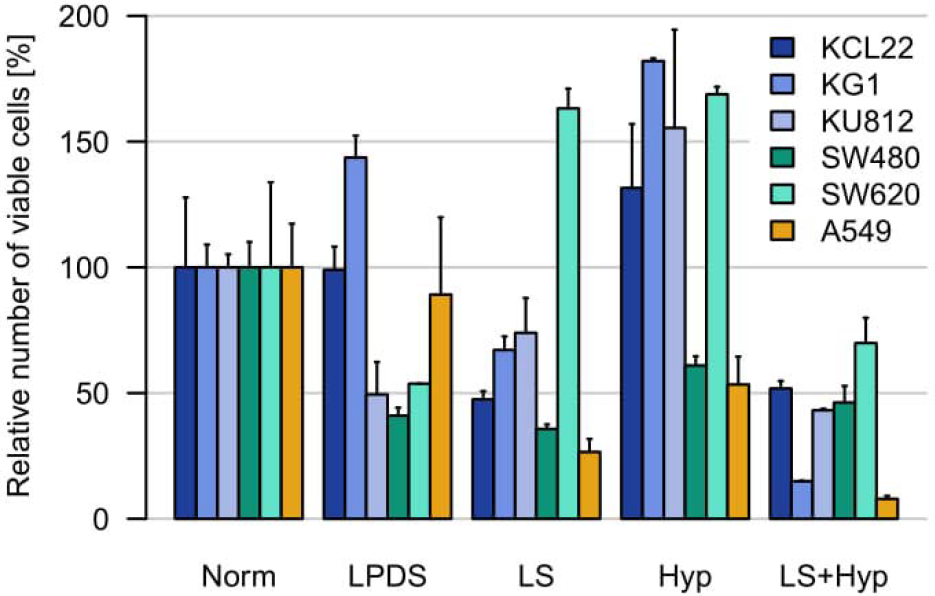
Effect of metabolic stress on proliferation rates different cancer cell lines. KG1, KCL22, KU812, SW480, SW620 and A549 cells were cultivated under lipoprotein deficient medium (LPDS serum), low-serum (LS) medium (2% serum), hypoxia (2% O2), or hypoxia in combination with LS medium. The number of live cells was counted using a trypan blue dye exclusion method, after 48 hours of culturing. To allow comparison between cell lines, numbers were normalized to control conditions.

It has been reported that cancer cell lines differentially regulate *de novo* lipid synthesis pathways under oxygen and/or nutrient deprivation (23-25, 28). Here, we also observed differential impact of metabolic stress on expression of FASN and HMGCR. This data also indicates cell-type specific regulation of *de novo* lipid synthesis pathways under metabolic stress (**Supplementary Figure 3**). mRNA levels of FASN and HMGCR under stress conditions also displayed highly positive correlation (R^2^=0.75; P <0.0001) (**Supplementary Figure 4**). The expression of markers of lipid uptake/degradation (LPL, MGLL and CD36) was also differentially affected by metabolic stress (**Supplementary Figure 3**).

### Effect of metabolic stress on lipidomic profile of cancer cells

Next, we studied the effects of metabolic stress on lipidomic profiles of cancer cells. Previous studies have already shown various stress-related effects on individual lipid-moieties. Our broad lipidomic assay allowed us to assess the impact of metabolic stress on robust averages of broader lipid-classes, providing a more holistic overview of cancer cell lipidomic profiles. **Figure 4** summarizes the data on lipidomic profiles of all cell lines cultivated under different metabolically-stressed conditions. We further stratified lipid classes on the basis of saturation index of their fatty-acyl side chains (expressed as the largest number of double bonds in any acyl side chain). *De novo* fatty acid synthesis enriches the cancer cell with saturated and/or mono-unsaturated fatty acids (20). Hence, selection of fatty acid acquisition mode –*de novo* FA synthesis versus FA uptake– by cancer cells affects saturation-index of fatty acyl side chains at least in membrane phospholipids (20). Here, we expressed the data relative to the baseline level (without stress) of the cell line. Thereby, we remove the existing baseline differences among cell lines as already discussed above for cell viability and thereby emphasize on changes in relative lipid distribution under stress. In each panel the data-points spreading away from the median-plane display differences from the normal conditions, color coded for lipid classes and stratified according to saturation index on the second dimension. We observed that cells cultivated under LPDS and LS containing media show multiple aberrations in their lipidomic profiles (**Figure 4**). Almost all cell lines showed significant decrease in cellular levels of CE under lipoprotein deficient conditions. Under LS medium all leukemic cells displayed decreased levels of CE of polyunsaturated fatty acyl chains. However, the cell lines derived from solid tumors displayed overall decrease in CE under serum-deprivation. Most striking alterations include changes in TG profiles of leukemia cell lines (KG1, KCL22 and KU812) cultivated under LS medium. In leukemia cells under serum-deprived conditions the cellular level of TG subspecies decreased with increasing number of double bonds in their fatty acyl chains. A similar but slightly less pronounced effect was observed in SW620 cells. In other words, LS medium induced decreased levels of polyunsaturated fatty acid-enriched TGs in selected cancer cell lines. Levels of DGs were increased in leukemia cell lines under LS conditions. Moreover, the levels of highly saturated PCPs were also significantly increased under LS condition particularly in KG1, KCL22 and KU812 cells. Previous studies have shown that tumor-associated lipogenesis increases the saturation of phospholipids, particularly PCs, in human cancer cells (20). Here, we did observe increased expression of FASN under LPDS medium in multiple cell lines. However, cellular levels of various PC subspecies did not display marked changes or specific trends under metabolic stress.

**Figure 4:**
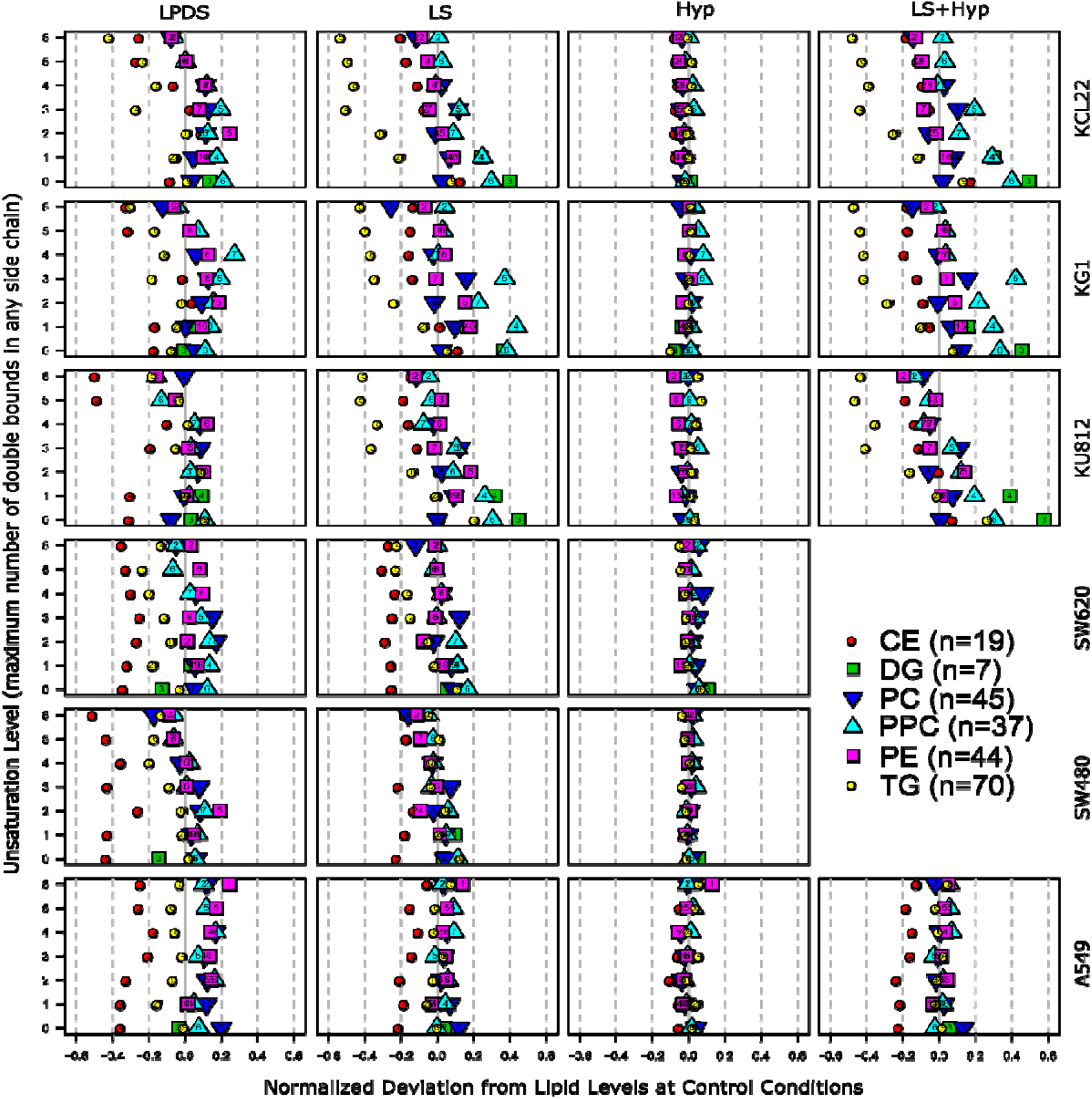
Effect of metabolic stress on lipidomic profiles of cancer cells: Each column shows changes in cellular levels of the six major lipid classes –including CEs, DGs, PCs PPCs, PEs and TG– under a specific stress condition relative to control for KCL22, KG1, KU812, SW620, SW480 and A549 cells. For each lipid class the peak intensities of the subspecies containing similar number of double bonds in their fatty acyl chains (chain containing highest number of double bonds) were summed up and the data were log2 transformed and median normalized. LS+Hyp stress was not tested for SW480 and SW620 cell lines.

Surprisingly, we did not observe any robust changes in lipidomic profiles of cancer cells cultivated under hypoxia. These results are strikingly clear for all of the detected lipid classes. Most of the previous studies have separately compared data for each lipid subspecies, whereas here we compared subgroups classified according to the number of double bonds in their fatty acyl chains. When individual lipid molecules were compared indeed certain changes in the cellular levels of various lipid subspecies could be observed. **Supplementary table 2** compares levels of various lipid subspecies in A549 cells cultivated under normoxia vs. hypoxia. Here, we also observed significant alterations in the levels of various lipid subspecies. Some of these changes were in line with published data (27, 31).

## Discussion

The presented work aimed to determine the impact of metabolic stress on expression of selected lipid metabolism genes, lipidomic profiles and cell proliferation in biologically diverse cancer cells. We observed that under metabolic stress different cancer cell lines showed differential expression of markers for *de novo* lipid synthesis and lipid-uptake/degradation. These data confirm previous works that showed cell-type specific regulation of lipid metabolism under oxygen and nutrient-deprivation (28). Careful scrutiny of literature revealed that previous works not only used different cell line models but also applied different cell culture conditions. For instance, differences in methods for inducing hypoxia or duration/severity of hypoxia were noted. These differences could have introduced variations in the data. Lewis CA *et al*. (32) have shown that U87 glioblastoma cells cultivated under hypoxia in full serum conditions display decreased expression of various lipid metabolism genes, in particular FASN and HMGCR. However, same cells when cultured under hypoxia in low-lipid conditions maintained high expression of these genes. Taken together these observations indicate that hypoxia and nutrient-deprivation may synergistically regulates lipid metabolism in cancer cells.

The major goal of the presented work was to study the impact of nutrient and oxygen-deprivation on lipidomic profiles of cancer cells. We observed that under low-lipid and low-serum environments the lipidomic profiles of cancer cells were significantly altered. Particularly, striking alterations in composition of neutral lipid (triglycerides and cholesterol esters) pools were noted. Previous studies have shown that cancer cells store excessive lipids and cholesterol in lipid droplets (LDs) (1, 33-35) – that are cytoplasmic deposit for triglycerides (TGs) and cholesterol esters (CE). Higher LDs and stored-cholesteryl ester (CE) content in tumors are now considered as hallmarks of cancer aggressiveness (1, 35-38). Moreover, LD-rich cancer cells are more resistant to chemotherapy (39). Cultured cells are shown to generate and increase the number of LDs in two different situations from a physiological perspective: when maintained in medium containing excess free fatty acids and/or lipoproteins or under nutrient- or oxygen-deprivation. Most of the previously published works have not focused on CE and TG subspecies profile in cancer cells. Here, we observed that under nutrient-deprivation cellular levels of all CE subspecies were significantly reduced, while the levels of TG containing polyunsaturated fatty acids were significantly reduced in all leukemia cell lines. Previous works on accumulation of LDs under stress utilized complete nutrient-deprivation (40). This severe stress leading to cell death induce the formation of LDs (41, 42) and in fact LD accumulation was considered to be a hallmark of apoptosis (43). In the presented work the cells cultivated under low-lipid or low-serum media were not under severe cytotoxic stress. Hence, it is possible that under mild stress cancer cells relied on previously stored LD content for the release of free fatty acids that subsequently caused decrease in CE and TG subspecies levels. It has been previously reported that under restricted cholesterol-rich low-density lipoprotein (LDL) supply cancer cells mobilize CEs (34). Cancer cells are shown to consume fatty acid (FA) through fatty acid β-oxidation (FAO) that is considered as the dominant bioenergetic pathway in non-glycolytic tumors. It has been shown that the dependence of cancer cells on FAO is further heightened in nutrient-deprived conditions (21).

The most striking observation of this study was that the lipidomic profiles of cancer cells remained robust under hypoxia. This effect was consistent throughout the selected cell line panel and for all the analyzed lipid sub-groups. This effect was particularly interesting because in line with previous works we also observed changes in proliferation capacity, expression of lipid synthesis/degradation markers and expression of hypoxia markers (data not shown) in hypoxic cancer cells. Despite all these changes cancer cells maintained homeostasis for all lipid classes. Also, if stresses are combined (LS+Hyp) the observed pattern is similar to LS conditions and not altered by hypoxia. Few previous studies reported that hypoxia induces changes in phospholipid/triglyceride profiles of cancer cells (26, 27, 31). Previous works compared the levels of individual lipid moieties and observed significant changes in multiple TG and PC subspecies. We also observed changes in cellular levels of various lipid subspecies when compared individually. However, our broad lipidomic assay allowed us to focus on robust averages from larger lipid-classes classified on the basis of saturation-density of their fatty acyl chains. These analyses provide a more holistic view of saturation status of various lipid classes. A previous study by Yu *et al*. (31) –that reported clear cut changes in PC profiles of Hela cells under hypoxia– also noted that changes in PC profiles are only evident when individual PC species are analyzed. However, when the relative abundance for PL species with acyl chains containing ≥3 double bonds were compared with that containing ≤3 double bonds no significant difference was observed between cells under normoxia and hypoxia. Meticulous literature survey revealed differences in cell culture methods among previous studies that may also lead to contradictory evidence. For instance; cells were serum-starved prior to hypoxia-induction (31), hypoxia was applied in combination with nutrient-deprivation (27) or media full serum media was supplemented with exogenous lipids (44). Here, we also observed that the lipidomic profiles of the selected cancer cell lines cultivated under low-serum environment were similar to that cultivated under hypoxia+low-serum environment. Hence, one can speculate the altered lipidomic profiles are mainly regulated by nutrient-availability and not by hypoxia.

## Acknowledgements

Work from the author’s laboratory was supported by Alexander von Humboldt Foundation and Higher Education Commission of Pakistan (Project # 2505/R&D/11-2670). We would like to thank Dr. Nadine Rohwer for help with determining expression of hypoxia markers in all samples.

## Abbreviations

FASN: fatty acid synthase
HMGCR: 3-hydroxy-3-methylglutaryl-CoA reductase
MGLL: monoacyl glycerol lipase
LPL: lipoprotein lipase
PC: Phosphatidylcholines
PE: Phosphatidylethanolamines
PPE: Plasmenylphosphatidylethanolamines
PPC: Plasmenylphosphatidylcholines
TG: Triacylglycerols
CE: Cholesterol esters
DG: Diacylglycerols

